# DYNAMITE: a phylogenetic tool for identification of dynamic transmission epicenters

**DOI:** 10.1101/2021.01.21.427647

**Authors:** Brittany Rife Magalis, Simone Marini, Marco Salemi, Mattia Prosperi

## Abstract

Molecular data analysis is invaluable in understanding the overall behavior of a rapidly spreading virus population when epidemiological surveillance is problematic. It is also particularly beneficial in describing subgroups within the population, often identified as clades within a phylogenetic tree, that represent individuals connected via direct transmission or transmission via differing risk factors in viral spread. However, transmission patterns or viral dynamics within these smaller groups should not be expected to exhibit homogeneous behavior over time. As such, standard phylogenetic approaches that identify clusters based on summary statistics (e.g., median genetic distance over the clade) would not be expected to capture dynamic clusters of transmission. For this purpose, we have developed DYNAMITE (DYNAMic Identification of Transmission Epicenters), a cluster identification algorithm based on a branch-wise (rather than traditional clade-wise) search for cluster criteria, allowing partial clades to be recognized as clusters. Using simulated viral outbreaks with varying cluster types and dynamics, we show that DYNAMITE is consistently more sensitive than existing tools in detecting both static and dynamic transmission clusters. DYNAMITE has been implemented in R and released as open source at: github.com/ProsperiLab/DYNAMITE.

## Introduction

Infectious disease epidemics are by nature dynamic, varying in size of the infected population, typically characterized by early explosive growth followed by either a decline that leads to extinction or an endemic steady state. Variation in infectious disease population dynamics can also be attributed to outside forces, such as public health interventions that would act to dampen the effects of disease spread. It has long been recognized that some individuals, or high-risk groups, in a population may transmit a pathogen more effectively than others because of specific biological, geographical, behavioral, or other “super-spreading” factors (e.g., [3, 12, 1]), thereby complicating predictions of epidemic growth or decline. Super-spreading individuals usually share common risk factors (e.g., health worker occupation) that can be identified and traced through traditional epidemiological surveillance techniques. Gathering such data, however, is often difficult, especially during the early stages of an epidemic when transmission routes might still be under validation, for example, or when resources limit contact tracing efforts. Molecular surveillance methods offer a rapid approach to identification of transmission clusters associated with these risk groups through linkage of sampled individuals based on pathogen genetic sequence data.

Numerous tools exist to identify transmission clusters using pathogen sequence data (e.g., reviewed in [7]). These tools are based on the assumption that direct transmission events can be observed in the sequence data when a maximum genetic distance threshold is set –i.e., patients with minimal sequence evolution are likely to have experienced less time between infection and thus fewer potential intermediate players. Multiple individuals can be connected to form a network, or cluster, of highly genetically similar samples. Network, or distance-based, tools such as HIV-TRACE [5] can also provide visualization of patients’ characteristics within clusters. Alternatively, phylogeny-based methods rely on the phylogenetic relationships among sequence data, (e.g., [8, 2, 10]. Minimal genetic distances define putative direct transmission events, similar to distance-based methods such as HIV-TRACE; however, distances are defined according to the branch lengths separating individual leaves within the tree (patristic distances) and clusters are typically required to be monophyletic clades (with the exception of PhyClip[2]) with a well-supported ancestral node. Support can be provided, for example, in the form of bootstrapping performed during tree reconstruction (e.g., at least 90% of bootstrappped trees)[4]. Overall, there has been good concordance among phylogeny- and distance-based methods in identification of clusters [13]. Yet, both methods rely on summary statistics of genetic distances (e.g., mean or median patristic distance within a putative cluster), assuming a relatively normal distribution of branch lengths within the clade and ignoring skewed variation as a result of the dynamic nature of transmission. For example, in a scenario of declining transmissions mediated by intervention targeting a particular risk group, the early spread within the cluster may be masked, or overwhelmed, by subsequent longer genetic distances separating delayed transmission if this pattern of transmission has occurred for a longer period of time than the period of growth. While there is a need to rapidly, and accurately, identify epicenters of transmission growth, relatively low-contributing transmission among groups of individuals in response to already implemented mitigation efforts are also of importance to future planning.

## Methods

### Cluster identification

Distinctly from other phylogeny-based clustering tools (e.g., Phylopart [8]), which consider only external leaves, DYNAMITE employs a branch-wise algorithm, which analyzes branch lengths among internal nodes, aiding in the identification of rapidly growing transmission clusters. Similar to Phylopart, an outlying fraction of the distribution of median patristic distances for each subtree within the full tree is considered as a cut-off, or threshold, representing the maximum genetic distance separating two nodes (internal or external) within the tree [8]. Whereas phylogeny-based tools such as Phylopart use this threshold to classify well-supported clades as clusters using summary statistics (e.g., median patristic distance within the clade), DYNAMITE’s branch-wise algorithm proceeds from the most recent common ancestor of the clade, accepting or rejecting subsequent branches (internal or external) within the tree according to the cut-off, allowing for partial clades to be categorized as transmission clusters. This is beneficial for two reasons: 1) a transmission cluster may be well-supported but harbor few highly divergent branches as a result of intense positive selection, false positive support, or the accumulation of sequencing errors, for example [2]; 2) a rapidly, or exponentially growing, cluster, whose branching pattern is characterized by shorter branches nearer the most recent common ancestor and long external branches, which would result in an overall larger median branch length. The algorithm works as follows (see Figure 1 for a graphical scheme): for each well-supported clade within the tree (provided by user according to support method of choice, such as bootstrapping), we begin at the ancestral node (considered level *L*_0_) and remove each branch (*b_j_*) in level *L*_*i*=1_ for which *b_j_* is greater than the branch length cutoff 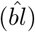 identified as described above. For each branch *b*_*j*(*i*+1)_ in subsequent level *L*_*i*+1_, mean branch length 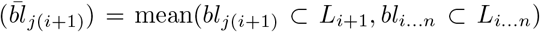. If 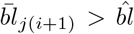, the node and corresponding subtree, or clade, is pruned. This is repeated for all subsequent levels until all external branches within the clade have been accepted or rejected. As downstream analyses require strictly bifurcating trees, parent nodes contributing to only one child in the cluster are cross-referenced against the original tree, and the corresponding child node is added back to the cluster.

**Figure 1:**
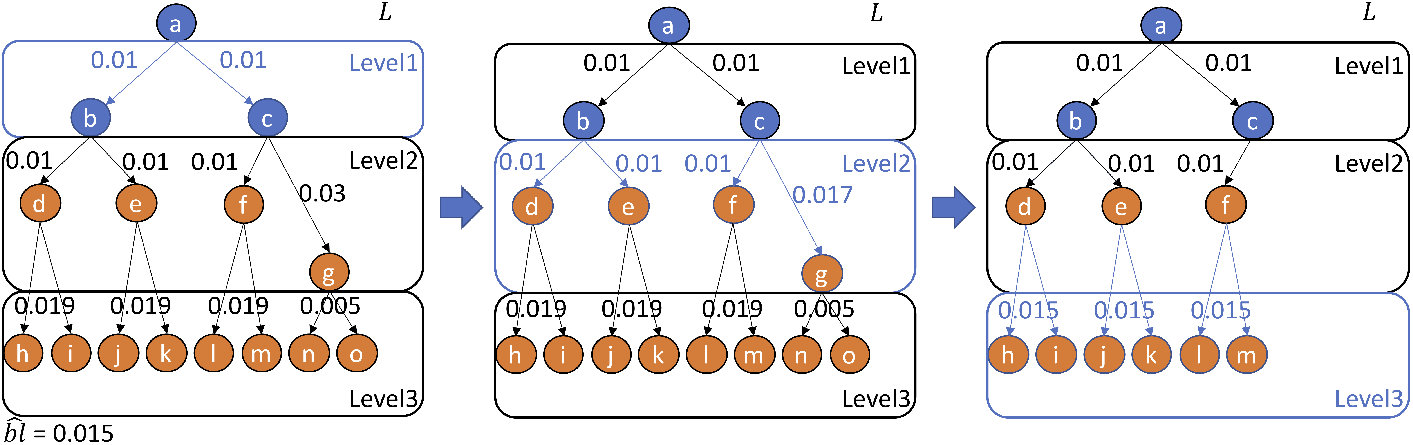
Schematic of DYNAMITE’s branchwise cluster-picking algorithm. Levels correspond to the generations of children nodes (b-o) descending from the original parental node (a), with nodes representing individual viral sequences derived from patient samples (h-o) or ancestral to patient samples (a-g). Edges, or branches, represent the relationship among nodes, with initial (black) values representing the genetic difference between immediately connected nodes. Updated (blue) values represent the mean of all branch lengths thus far on the reverse path to the most recent common ancestor (a), with each level evaluated separately, as described in the Methods. Edges (and downstream nodes) that result in an updated mean branch length ¿ pre-specified branch length cutoff 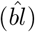 are removed (e.g., (g,(n,o))).

### Simulation of transmission clusters

Simulation of epidemic outbreaks was performed using the nosoi [6] agent-based stochastic simulation platform. Seven distinct sub-populations [B-H] were allowed to emerge from the background population (A) with a probability of initial infection of 7.5 E — 04 after at least two background individuals had been infected at the start of the simulation. Following initiation, transmission was isolated to the sub-population (i.e., probability of zero of infecting an individual in another group), with the exception of subgroup H, which was the sole contributor to infection in the eighth sub-population, I (probability of infection 1.5*E* – 02). The number of contacts for groups A-E and H-I were picked from a normal distribution with group-specific means and standard deviation of one (Table 1). The number of contacts for E and F, however, were derived from the following function:

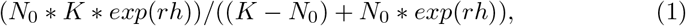

where *N*_0_ is the initial number of contacts, *K* is the maximum, and *r* is the rate of change dependent on the current number of actively infected hosts in the simulation (h). Sub-population F, with a positive *r* was considered to be growing at a more rapid rate than remaining sub-populations, whereas G, with a negative r was considered to be decaying (Table 1). The probability of transmission (when a contact occurs) was provided in the form of a threshold function: before a certain amount of time since initial infection, the host does not transmit (mean incubation time of 5 days (sd=2)), and after that time, the individual will transmit with a certain (constant) probability (Table 1). I.e., this function is dependent on the time since the host’s infection. Probability of transmission was also dependent on subgroup, resulting in a range of basic reproductive numbers (*R*_0_) for each subgroup differing according to number of contacts, probability of transmission, or both. For example, The *R*_0_ for subgroups C and D were both 5.5, but these two groups differed in transmission-related parameters. This was designed to test whether branching patterns would differ depending on the individual parameters comprising *R*_0_, thus influencing cluster identification.

**Table 1:**
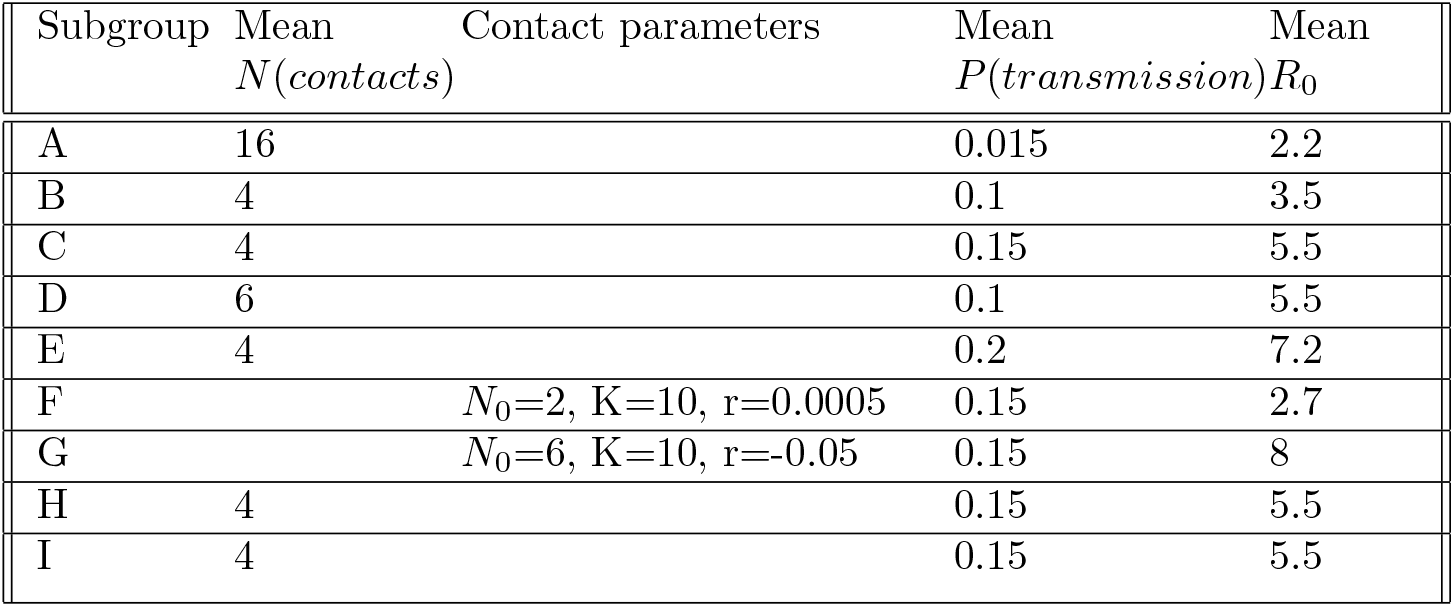
Simulation information for each subgroup.

Each of a total of 1,000 simulations was run for 365 days or until a total of 10,000 hosts were infected. One representative clade within the full phylogenetic transmission tree for each subgroup (including the background population) was chosen at random from the corresponding internal nodes that contained between 5 and 30 external taxa, representing true clusters of direct transmission (e.g., Figure 2). A random sample for a total of 3X the length of the simulation of the remaining population was combined with the true clusters. Hosts not included within this sample were pruned from the full tree to obtain the final 1,000 simulated trees used for transmission cluster identification. A molecular clock, or constant evolutionary rate across all branches of the tree, was assumed, allowing branches separating noes within the tree to be scaled in both time and genetic distance.

**Figure 2:**
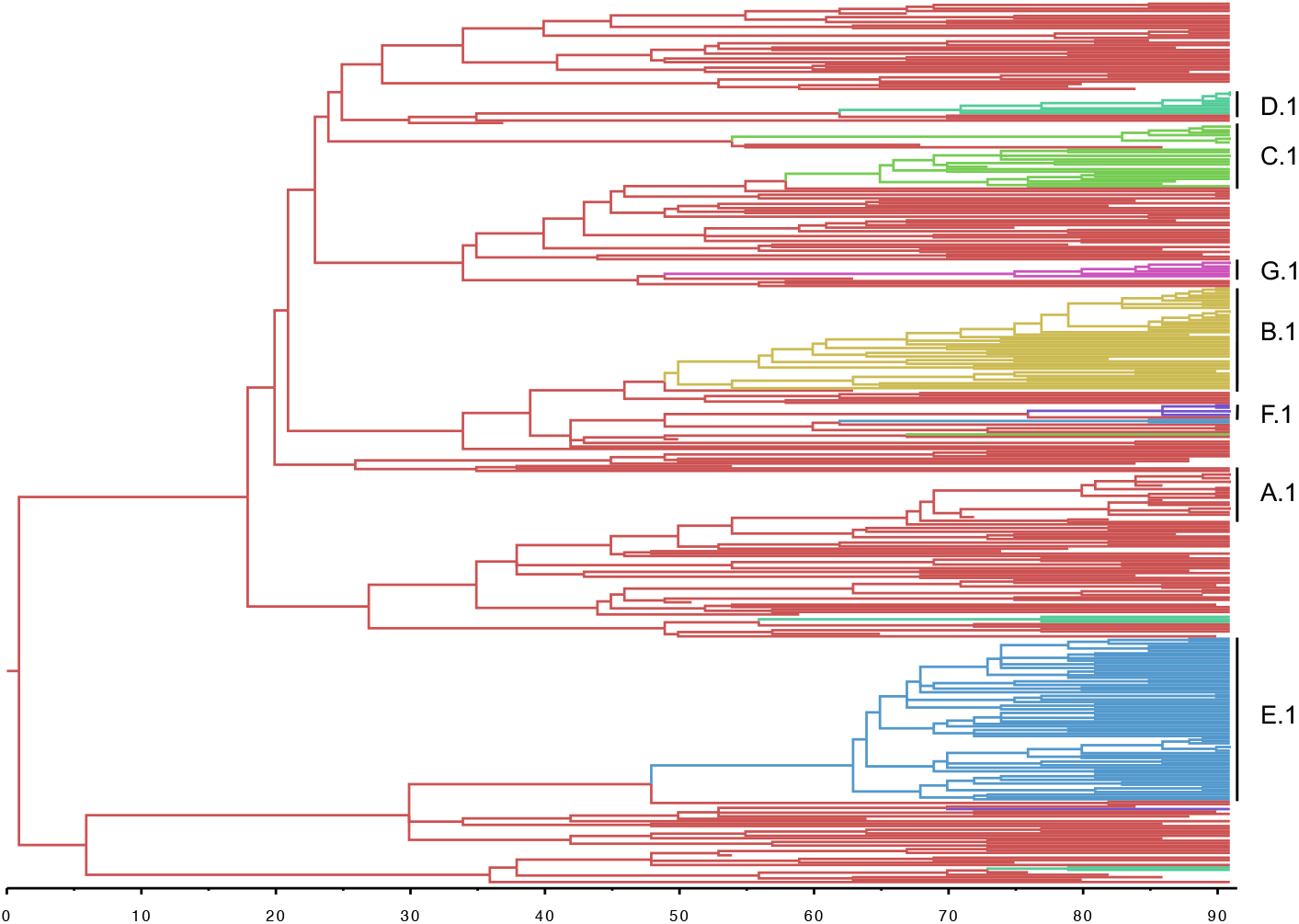
Example tree connecting randomly sampled infected individuals from the background (red) population and individuals linked via direct transmission through varying transmission dynamics (A-G). Individuals from subgroups H and I described in the Methods section did not emerge in this (second) simulated epidemic.

### Evaluation of performance

A range of branch length thresholds from 1-20% was used to evaluate the ability of DYNAMITE’s branch-wise algorithm to identify the various types of transmission clusters described above, as well as the that of the more traditional clade-wise algorithm of Phylopart [8]. Node groups or clades classified as transmission clusters by the branch-wise or clade-wise algorithm, respectively, were considered true clusters if at least 70% of the true taxa were contained within the identified cluster.

## Results

After simulating clusters of differing starting transmission potential and transmission dynamics over time, performance of DYNAMITE’s branch-wise algorithm was compared to the standard clade-wise approach of Phylopart [8], as well as for individual cluster subgroups (A-F). The overall ability of the branchwise algorithm to identify clusters was measured in terms of the fraction of true positive clusters identified (TPR) and fraction of all clusters identified that were not true positive (FPR) (Figure 3). Whereas the clade-wise algorithm begins to plateau at a median ratio of TPR to FPR of approximately two, the branch-wise algorithm continues to climb, reaching a median TPR/FPR of four at the same final branch length threshold of 20%.

**Figure 3:**
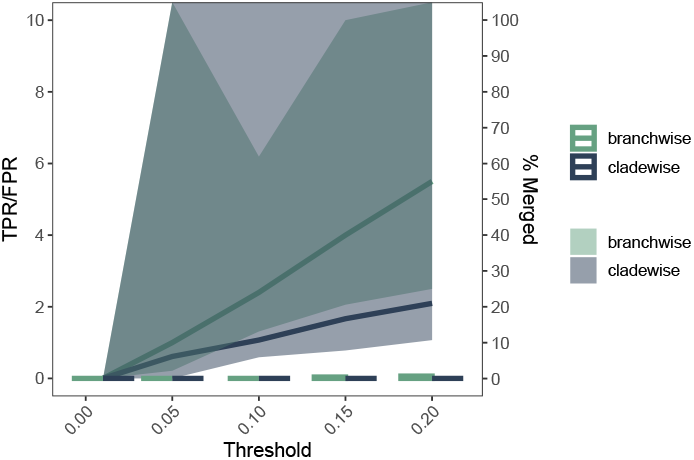
Rate of detection of true transmission clusters using the clade-wise (i.e., Phylopart) and branch-wise algorithms. True positive rate (TPR) is expressed as the percentage of true clusters identified using each algorithm. False positive rate (FPR) is expressed as the percentage of all clusters identified that are not considered to be true clusters. The median ratio (TPR/FPR) and 95th percentile ranges are represented by solid line and surrounding shaded areas (y-axis, left). Bars represent the percentage of identified clusters that were identified as a single cluster (y-axis, right).

As true positive clusters were identified according to the percentage of true taxa, the possibility of merging several phylogenetically close clusters into one single cluster grows with increasing branch-length thresholds. A comparison of the percentage of true clusters merged into a single cluster was made between the two algorithms, but the frequency of this occurrence was low for both (Figure 3).

Subgroup A is characterized by an *R*_0_ (2.2) of that of the background, or majority, population, and thus represents a contact-traced subgroup with the same risk for transmission as the majority of infected individuals. The ability to detect direct transmission clusters is especially critical in the early part of an epidemic when the risk factors for spread are as-of-yet unidentified. The branchwise algorithm outperforms the clade-wise one in the detection of subgroup A alone (Figure 4), with a consistently one-fold greater TPR/FPR past the 5% branch length cutoff. However, peak TPR/FPR for both algorithms are lower for subgroup A (Figure 3) than overall (Figure 4, suggesting a bias toward smaller genetic distances that would be characteristic of, for example, higher transmission rates.

**Figure 4:**
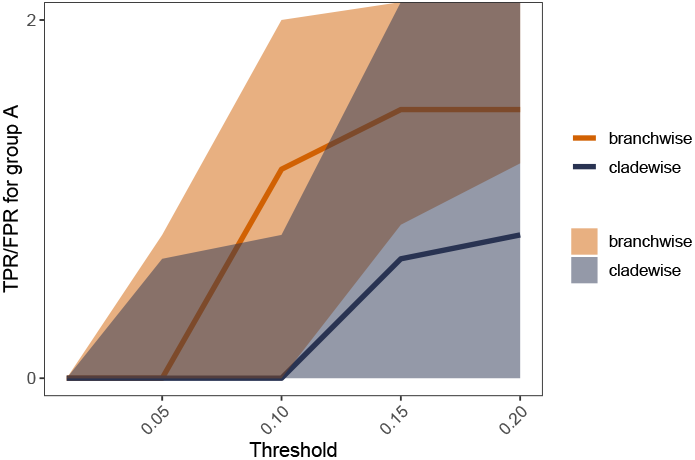
Rate of detection of true transmission clusters belonging to the background (Ro 2.2) population using the clade-wise (i.e., Phylopart) and branchwise algorithms. True positive rate (TPR) is expressed as the percentage of true clusters identified using each algorithm. False positive rate (FPR) is expressed as the percentage of all clusters identified that are not considered to be true. Median TPR/FPR and 95th percentile ranges are represented by solid line and surrounding shaded areas (y-axis, left)

Indeed, the majority of higher-Ro clusters were readily detected by both algorithms (red, Figure 5). Due to the stochastic nature of the simulation used for epidemic growth, it was possible for more than one of the same subgroup (B-I) to emerge during a single simulation. As only one of each subgroup present was chosen to represent an epidemiologically linked, or true, transmission cluster, any remaining subgroups were still present but at the sample sampling frequency as the background population (2-5%). These clades still represent high-risk groups, though not related in the tree via direct transmission. As many as 50% of identified clusters during a simulation belonged to this category, with the clade-wise algorithm identifying more than the branch-wise (blue, Figure 5). This is a key find, as it demonstrates that sampling of every individual involved in a high-risk transmission group is not necessary for the identification and characterization of these groups for which additional clinical data would aid in improving more targeted interventions.

**Figure 5:**
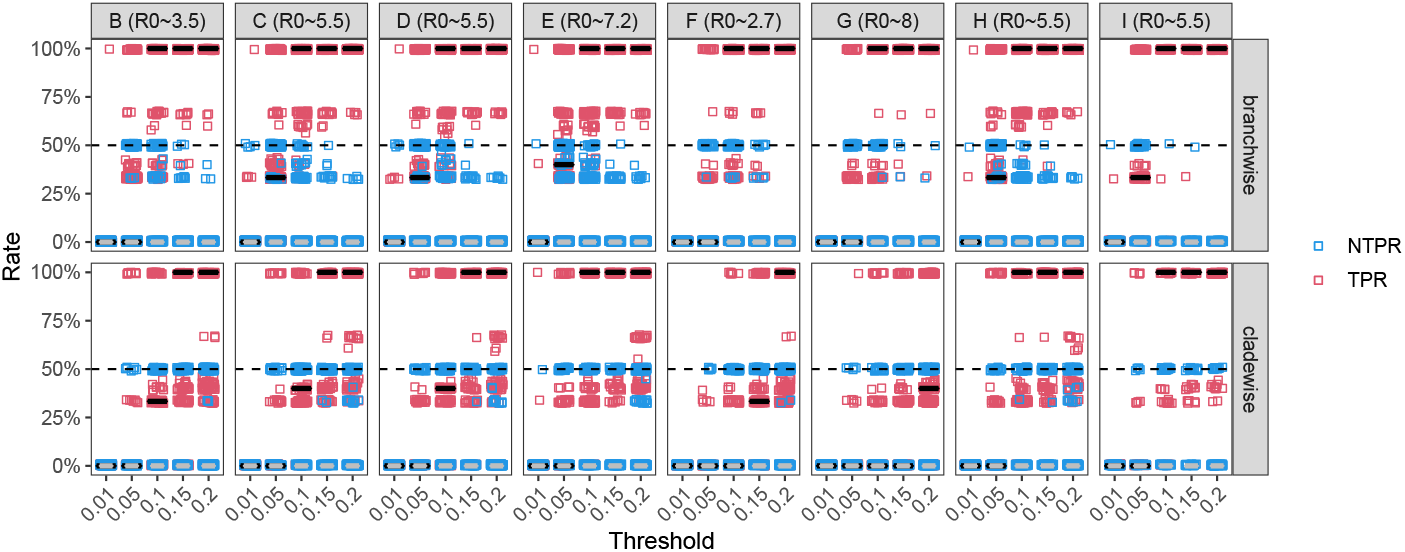
Rate of detection of true transmission clusters belonging to varying elevated Ro subpopulations using the clade-wise (i.e., Phylopart) and branchwise algorithms. Each point represents a single simulation. True positive rate (TPR) is expressed as the percentage of true clusters identified using each algorithm. Negated true positive rate (NTPR) is expressed as the percentage of all clusters identified that are not considered to be true (i.e., truly connected via direct transmission) but still high risk groups.

Whereas the majority of static high-transmission clusters were identified by the clade-wise algorithm (Figure 5, the ability to detect the rapidly growing cluster (F) with a median TPR comparable to the branch-wise algorithm was limited to the highest threshold (20%), and the cluster characterized by declining transmission potential (G) was only detected at a median TPR of approximately 30% at the highest threshold. This scenario is problematic as the FPR increases more rapidly after a threshold of 10%, resulting in a high-risk, low-reward trade-off between the identification of dynamic clusters and false positive clusters.

Understanding the temporal dynamics of a cluster over time, such as the distribution of individuals with specific risk factors, as well other temporal characteristics, such as timespan and origin, critically depend on good coverage over the time period of the existence of the cluster. As DYNAMITE’s branch-wise algorithm relies heavily on the early branches within a clade, we sought to determine if the step necessary to force bifurcation whereby children nodes are added to non-bifurcating parents was sufficient to extract this necessary information from the identified cluster (Figure 6). Indeed, the median proportion of true taxa identified by the branch-wise algorithm for a threshold ¿ 5% was one, rivaling that of the clade-wise algorithm.

**Figure 6:**
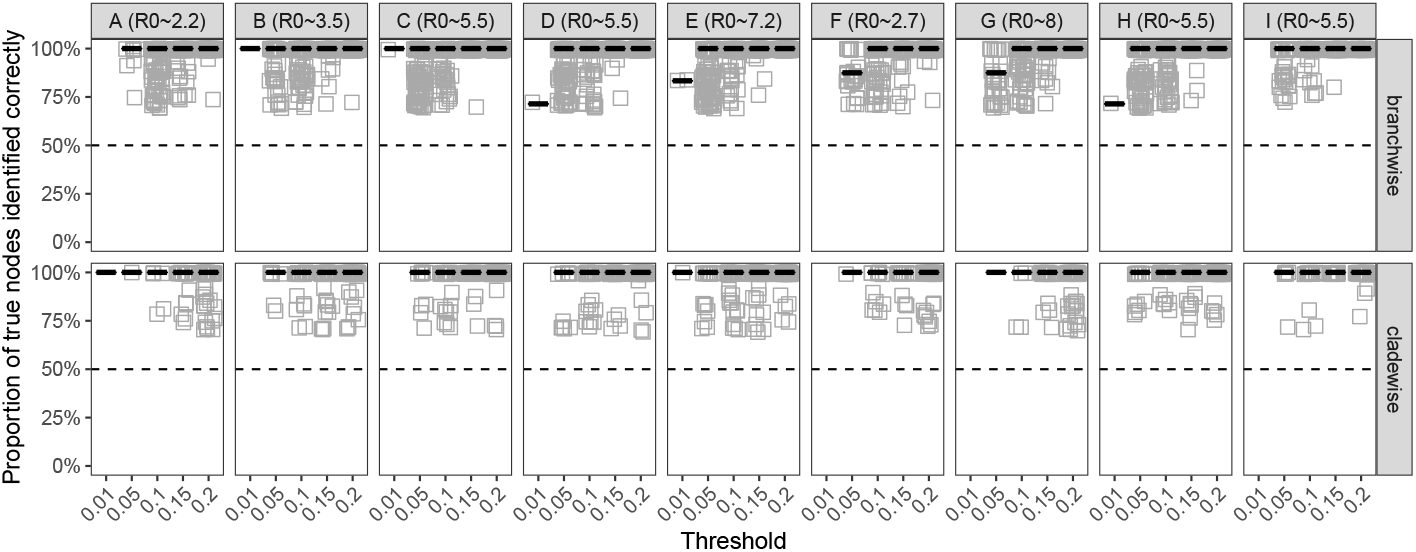
Proportion of nodes belonging to varying elevated *R*_0_ transmission clusters identified using the cladewise (i.e., Phylopart) and branchwise algorithms. The proportion of phylogenetic nodes (internal and external) identified is expressed as a percentage of the total nodes belonging to the true transmission cluster.

The DYNAMITE code, implemented in R v¿3.6 [9] is available from github (github.com/ProsperiLab/DYNAMITE), as are the R scripts used for simulation and benchmarking, and resulting simulated trees. For a single tree with 362 tips (Figure 2), the DYNAMITE branch-wise algorithm completed in 5.571 seconds in R v4.0 on a MacBook Air (2 GHz Intel Core i7, 8GB). DYNAMITE thus provides a scalable approach to identifying dynamic clusters for large outbreaks.

## Discussion

When compared to its phylogenetic ancestor, Phylopart [8], DYNAMITE’s branch-wise algorithm identified with greater accuracy not only direct transmission clusters belonging to the background population (equivalent transmission rates) and high-risk populations (higher transmission rates), but also dynamic clusters characterized by growth or decline. The branch-wise algorithm uses the initial Phylopart step of relying on patristic distances, or branch lengths separating external leaves, within the entire tree to determine the threshold criterion for cluster identification and proceeds to find well-supported nodes. Diverging from Phylopart and other other phylogeny-based approaches, each subsequent branch (internal or external) stemming from a well-supported node is considered a separate entity that is either accepted or rejected based on its relationship to the determined cutoff value. The concept of rejecting unusually long branches within a putative cluster has also been introduced recently in PhyClip [2], as highly divergent branches in well-supported clades can occur as a result of intense positive selection, false positive support, or simply the accumulation of neglected sequencing errors, for example. However, PhyClip, like Phylopart, relies solely on patristic distances, rather than examining internal branches individually. DYNAMITE’s branch-wise approach enables the identification of clusters for which the branch length distribution within the clade is highly skewed as a result of dynamic transmission patterns. These transmission patterns can include rapid growth or even decline, which was simulated according to an exponentially increasing or decreasing number of contacts for an infected individual in this study, though testing of additional dynamic patterns in the future might be warranted. This ability renders DYNAMITE particularly useful for upcoming SARS-CoV-2 epidemiological analysis; as vaccines become increasingly available, monitoring the impact of vaccination on transmission among particular risk groups will be necessary to determine further dissemination strategies in the upcoming year.

In addition to dynamic clusters, high-risk clusters were defined in this study as groups of individuals with elevated initial secondary infection rates, or basic reproductive number (Ro), and represent subgroups within the population associated with phenotypic factors that put the individuals at higher risk of infection. High-risk clusters not connected by direct transmission, whether static or dynamic, were detected at frequencies of nearly 75% by the branch-wise algorithm. Additional analyses to determine at what sampling fraction these subgroup can be identified are certainly warranted, as contact tracing is not always available for epidemiological analysis, and so high-risk populations are not always fully present in the phylogenetic tree. If unknown high-risk populations sampled at a slightly higher frequency than the background population can be detected and ruled out as non-contact-traced individuals, their associated clusters can be used to identify groups for further study on targeted intervention.

While we do not describe a way to discriminate high-risk clusters from direct transmission clusters, or dynamic from static clusters, or even a way to characterize cluster dynamics, approaches to do just this are under development and promise a way to better understand the inherently dynamic nature of transmission among individuals in real-world epidemics. A further limitation to DYNAMITE, like other phylogeny-based approaches, is the seemingly arbitrary threshold for cluster identification. Whereas in the current study, 10% of the whole-tree patristic distance distribution was sufficient to detect 100% of clusters with a low false positive rate, this may differ from dataset to dataset, depending on, for example, the expected number of clusters and sampling of the population. The maximum genetic distance separating direct transmission events is well-characterized for HIV based on the level of viral diversity reached within the host during infection [5]. This value (0.15 substitutions/site) and others can be used as a cutoff for most genetic clustering approaches; however, the intra-host diversity or similar information on direct transmission is not always known for emerging epidemics. For this reason, we encourage testing a range of thresholds surrounding 10% for comparison. But because DYNAMITE is not dependent on a pre-specified genetic distance cutoff, it can be applied to virtually any viral outbreak for which the virus is measurably evolving [11]. Similar to HIV-TRACE, DYNAMITE can also use metadata information supplied in the form of a taxa-identified table to pair with cluster data so that indexing is not required and cluster-related risk factors can be assessed more readily. DYNAMITE is thus a flexible tool applicable to all measurably evolving viruses that can be used to identify otherwise missed dynamic clusters which may be useful in public health intervention.

